# Comparing adaptive coding of reward in bipolar I disorder and schizophrenia

**DOI:** 10.1101/2021.10.24.465471

**Authors:** Mariia Kaliuzhna, Matthias Kirschner, Philippe N. Tobler, Stefan Kaiser

## Abstract

**Background:** Deficits in neural processing of reward have been described in both bipolar disorder (BD) and schizophrenia (SZ), but it remains unclear to what extent these deficits are caused by similar mechanisms. Efficient reward processing relies on adaptive coding which allows representing large input spans by limited neuronal encoding ranges. Deficits in adaptive coding of reward have previously been observed across the SZ spectrum and correlated with total symptom severity. In the present work we sought to establish whether adaptive coding is similarly affected in patients with BD.

**Methods:** 25 patients with BD, 27 patients with SZ and 25 healthy controls performed a variant of the Monetary Incentive Delay task during functional magnetic resonance imaging in two reward range conditions.

**Results:** Adaptive coding was impaired in BD and SZ in the posterior part of the right caudate. In contrast, BD did not show impaired adaptive coding in the anterior caudate and right precentral gyrus/insula, where SZ showed deficits compared to healthy controls.

**Conclusions:** BD patients show adaptive coding deficits, that are similar to those observed in SZ in the right posterior caudate. Adaptive coding in BD appeared more preserved as compared to SZ participants especially in the more anterior part of the right caudate and to a lesser extent also in the right precentral gyrus. Thus, dysfunctional adaptive coding could constitute a fundamental deficit in severe mental illnesses that extends beyond the schizophrenia spectrum.

## Introduction

Recent dimensional approaches to psychopathology highlight the continuity between schizophrenia (SZ) and bipolar disorder (BD) at different levels including clinical presentation, brain morphology, genetic markers and brain connectivity [1]. Alterations in reward processing have been described in both disorders, but previous literature remains inconclusive concerning continuity in reward processing deficits between the two disorders [2-5]. While in schizophrenia findings converge towards reduced striatal activation during the processing of reward, this result is less clear in BD. Thus, depending on the task, task stage (anticipation of reward or reward outcome) and patients’ clinical state (depressed, euthymic, manic) increased [6-8], decreased [4, 9-12] and similar [3, 13, 14] brain responses as compared to control participants have been observed in BD.

One of the core mechanisms underlying efficient reward processing in healthy participants is adaptive coding. Adaptive coding is the ability of neurons to adjust their response according to the present context [15, 16]. Extensively described in perception, adaptive coding is also characteristic of midbrain dopaminergic neurons, which, rather than coding the absolute value of a received reward, adapt their responses to the range of most probable rewards at a given time [17]. Physiological studies in non-human primates suggest that range adaptation allows for a more precise encoding of the stimulus, by making optimal use of the entire firing range of the neuron. In humans, several reward sensitive regions, such as the striatum, have been shown to adapt their response depending on the reward context [18]. Specifically, when the range of possible rewards is narrow (e.g. you can win up to 40 cents) these regions show a steeper response curve, that is, the BOLD response increases more strongly to increasing reward amounts received by healthy participants (Figure 1) [19, 20]. Conversely, when the range of possible outcomes is wide (e.g. you can win up to 2$) and more potential rewards need to be encoded, the increase in the BOLD response to increasing reward amounts is shallower.

**Figure 1.**
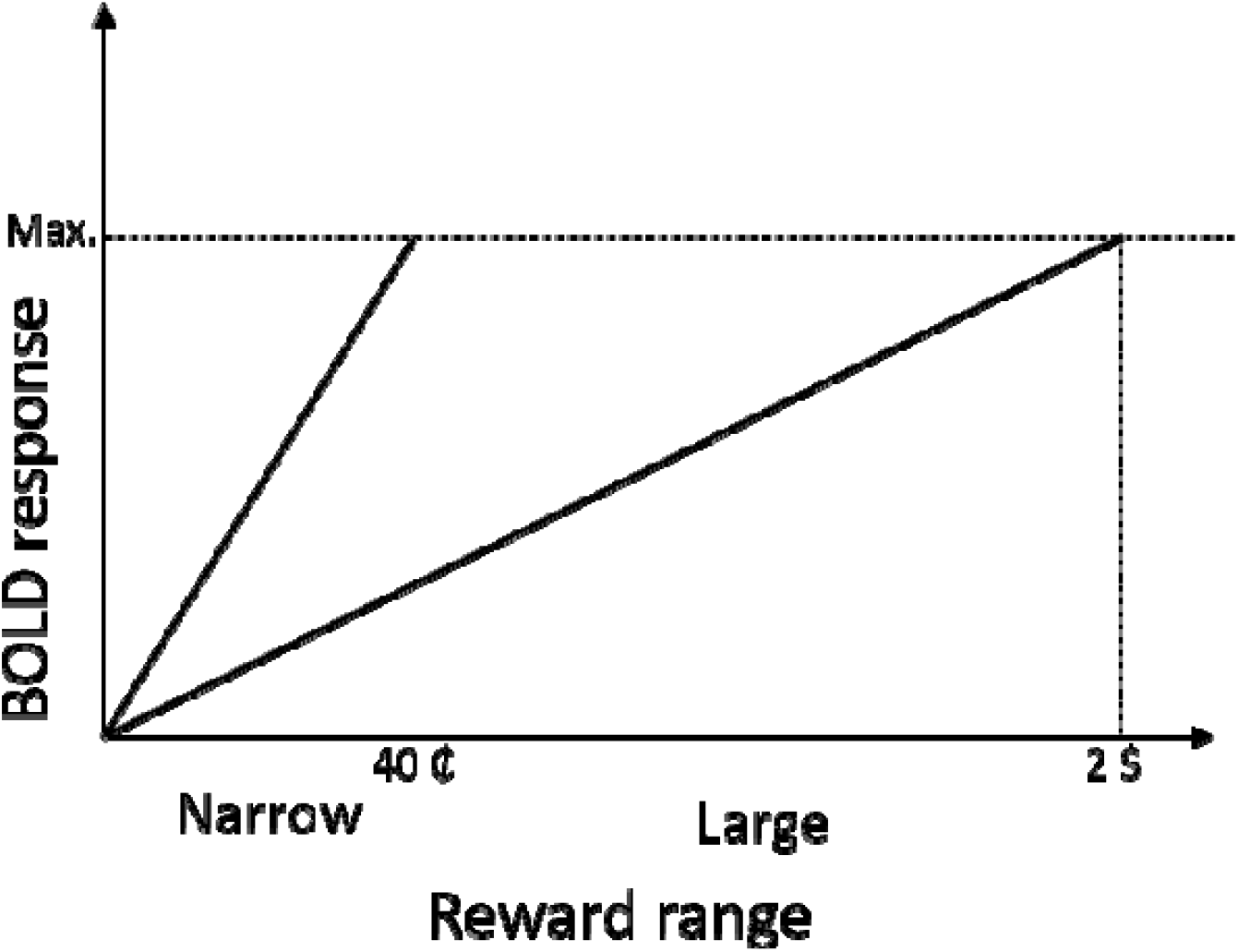
Schematic representation of adaptive reward coding during the MID task, as observed in the caudate of healthy controls. Efficient encoding of all possible rewards with a limited coding range requires the brain to dynamically adapt the response sensitivity to the currently available reward range. Accordingly, a more shallow slope of the BOLD response is expected (and observed) in the large reward range condition (2$) than in the small reward range condition (40⍰). By extension, the same reward (for example, 40⍰) will elicit different responses depending on the context it is presented in: a maximal response in the small reward range and an intermediate response in the large reward range condition.

Deficient adaptive coding of reward in the caudate and the insula/precentral gyrus has been described for the whole SZ spectrum, including participants with schizotypal personality traits, patients with first episode psychosis, as well as patients with schizophrenia [19, 20]. These participants show similar response increases for wide and narrow reward ranges, indicating reduced contextual influence. Although a dopamine dysfunction affects both SZ and BD [21], nothing is known about whether patients with BD can represent reward in an adaptive fashion.

In the present work we sought to establish whether adaptive coding of reward is reduced in euthymic patients with BD similarly to patients with SZ.

Here we used the same paradigm and analysis approach as in our previous work on adaptive coding of reward across the schizophrenia spectrum [19, 20]. Combining a version of the monetary incentive delay task with fMRI, we compare adaptive coding between three groups: BD patients, SZ patients and healthy controls. We sought to establish whether adaptive coding deficits are present in BD to a similar extent and in the same brain regions as in SZ, and, if the deficits, should they exist, relate to shared symptoms of BD and SZ.

## Methods and materials

### Participants

Twenty-five patients with BD-I participated in the study; 18 outpatients and 7 inpatients at the end of hospitalization (one patient was excluded from the analysis due to corrupt data files). Data from patients with schizophrenia and healthy controls have previously been reported by Kirschner and colleagues (2016), and included 27 patients with schizophrenia (16 inpatients and 11 outpatients) and 25 healthy controls (HC) (Table 1).

**Table 1.**
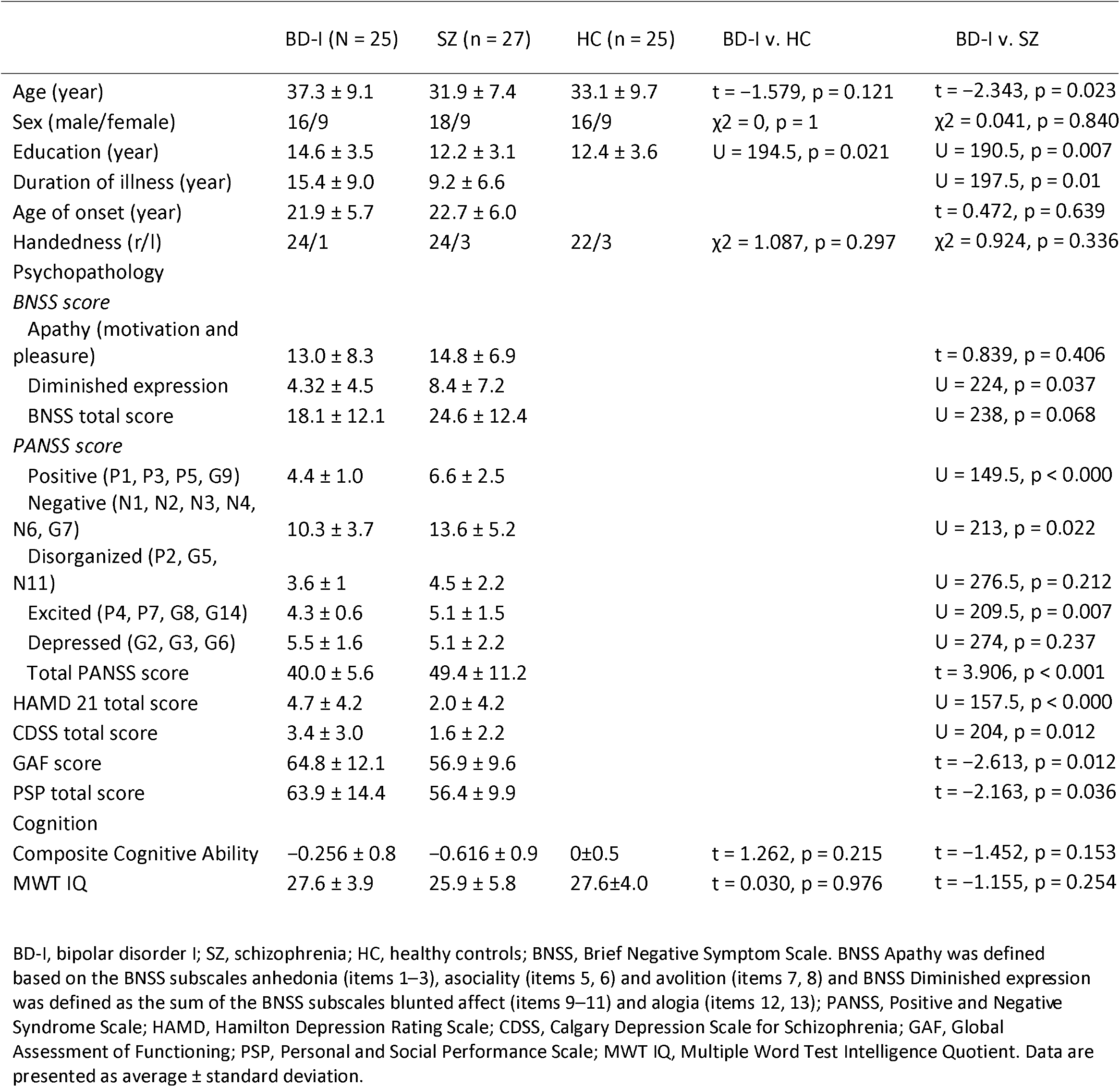
Socio-demographic and neuropsychological characteristics of the participants

|  | BD-I (N = 25) | SZ (n = 27) | HC (n = 25) | BD-I v. HC | BD-I v. SZ |
| --- | --- | --- | --- | --- | --- |
| Age (year) | 37.3 ± 9.1 | 31.9 ± 7.4 | 33.1 ± 9.7 | t = -1.579, p = 0.121 | t = -2.343, p = 0.023 |
| Sex (male/female) | 16/9 | 18/9 | 16/9 | χ<sup>2</sup> = 0, p = 1 | χ<sup>2</sup> = 0.041, p = 0.840 |
| Education (year) | 14.6 ± 3.5 | 12.2 ± 3.1 | 12.4 ± 3.6 | U = 194.5, p = 0.021 | U = 190.5, p = 0.007 |
| Duration of illness (year) | 15.4 ± 9.0 | 9.2 ± 6.6 |  |  | U = 197.5, p = 0.01 |
| Age of onset (year) | 21.9 ± 5.7 | 22.7 ± 6.0 |  |  | t = 0.472, p = 0.639 |
| Handedness (r/l) | 24/1 | 24/3 | 22/3 | χ<sup>2</sup> = 1.087, p = 0.297 | χ<sup>2</sup> = 0.924, p = 0.336 |
| **Psychopathology** |  |  |  |  |  |
| *BNSS score* |  |  |  |  |  |
| Apathy (motivation and pleasure) | 13.0 ± 8.3 | 14.8 ± 6.9 |  |  | t = 0.839, p = 0.406 |
| Diminished expression | 4.32 ± 4.5 | 8.4 ± 7.2 |  |  | U = 224, p = 0.037 |
| BNSS total score | 18.1 ± 12.1 | 24.6 ± 12.4 |  |  | U = 238, p = 0.068 |
| *PANSS score* |  |  |  |  |  |
| Positive (P1, P3, P5, G9) | 4.4 ± 1.0 | 6.6 ± 2.5 |  |  | U = 149.5, p < 0.000 |
| Negative (N1, N2, N3, N4, N6, G7) | 10.3 ± 3.7 | 13.6 ± 5.2 |  |  | U = 213, p = 0.022 |
| Disorganized (P2, G5, N11) | 3.6 ± 1 | 4.5 ± 2.2 |  |  | U = 276.5, p = 0.212 |
| Excited (P4, P7, G8, G14) | 4.3 ± 0.6 | 5.1 ± 1.5 |  |  | U = 209.5, p = 0.007 |
| Depressed (G2, G3, G6) | 5.5 ± 1.6 | 5.1 ± 2.2 |  |  | U = 274, p = 0.237 |
| Total PANSS score | 40.0 ± 5.6 | 49.4 ± 11.2 |  |  | t = 3.906, p < 0.001 |
| HAMD 21 total score | 4.7 ± 4.2 | 2.0 ± 4.2 |  |  | U = 157.5, p < 0.000 |
| CDSS total score | 3.4 ± 3.0 | 1.6 ± 2.2 |  |  | U = 204, p = 0.012 |
| GAF score | 64.8 ± 12.1 | 56.9 ± 9.6 |  |  | t = -2.613, p = 0.012 |
| PSP total score | 63.9 ± 14.4 | 56.4 ± 9.9 |  |  | t = -2.163, p = 0.036 |
| **Cognition** |  |  |  |  |  |
| Composite Cognitive Ability | -0.256 ± 0.8 | -0.616 ± 0.9 | 0 ± 0.5 | t = 1.262, p = 0.215 | t = -1.452, p = 0.153 |
| MWT IQ | 27.6 ± 3.9 | 25.9 ± 5.8 | 27.6 ± 4.0 | t = 0.030, p = 0.976 | t = -1.155, p = 0.254 |

BD-I, bipolar disorder I; SZ, schizophrenia; HC, healthy controls; BNSS, Brief Negative Symptom Scale. BNSS Apathy was defined based on the BNSS subscales anhedonia (items 1–3), asociality (items 5, 6) and avolition (items 7, 8) and BNSS Diminished expression was defined as the sum of the BNSS subscales blunted affect (items 9–11) and alogia (items 12, 13); PANSS, Positive and Negative Syndrome Scale; HAMD, Hamilton Depression Rating Scale; CDSS, Calgary Depression Scale for Schizophrenia; GAF, Global Assessment of Functioning; PSP, Personal and Social Performance Scale; MWT IQ, Multiple Word Test Intelligence Quotient. Data are presented as average ± standard deviation.

All participants were between 18 and 55 years old. The diagnosis of BD-I and schizophrenia was confirmed using the Mini-International Neuropsychiatric Interview for DSM-IV (MINI). We excluded patients with schizoaffective disorder, current major depressive, manic or hypomanic episode, as well as patients with any other DSM-IV axis I disorder or neurological disorder. All patients were clinically stable with no change to medication for at least 2 weeks prior to inclusion in the study. The absence of major extrapyramidal symptoms was confirmed using the Modified Simpson-Angus Scale (total score ≤2). BD-I patients were clinically euthymic and did not present with either more than sub-syndromal depressive symptoms (Hamilton Depression Rating Scale (HAMD) score < 17), as defined by the International Society for Bipolar Disorder Task Force [22] or manic symptoms (confirmed using the Young Mania Rating Scale [23], mean=0.4, sd=0.91, min=0, max=4). In HC the MINI was used to confirm the absence of any current or previous neurological or psychiatric conditions. The study was approved by the local ethics committee of the canton of Zurich and all participants provided written informed consent.

To clinically assess depressive symptoms we used the Hamilton Depression Rating Scale (HAMD 21 [24] and the Calgary Depression Scale for Schizophrenia (CDSS, [25]. Negative symptoms were assessed using the Brief Negative Symptom Scale (BNSS, [26] and factor scores for apathy and diminished expression were calculated according to Mucci and colleagues [27]. To assess the general level of functioning we used the Global Assessment of Functioning Scale (GAF, [28] and the Personal and Social Performance Scale (PSP, [29]. Psychotic symptoms were assessed using the Positive and Negative Syndrome Scale (PANSS, [30]. Cognition was assessed using the Brief Neurocognitive Assessment [31].

### Procedure

Participants performed a variant of the Monetary Incentive Daly Task (MID), originally developed by Simon and colleagues [32], that uses stimuli based on the Cued-Reinforcement Reaction Time Task [33]. Before the beginning of the task participants were informed that they would receive all the money they would win during the experiment. At the start of every trial one of three cues, signaling no reward, small reward, or large reward, indicated the reward context. In the no reward condition participants won nothing. In the small reward condition, they could win between 0 and 40 cents, and in the large reward condition they could win between 0 and 2 Swiss francs. The exact amount won depended on participants’ speed (RT on the current trial compared to RTs in the previous 15 trials) and accuracy (incorrect or late (>1s) response resulted in a reward of 0). The response was elicited by three circular shapes presented after the cue and the participants had to indicate the odd one out (right or left button press), responding as fast and as correctly as possible. Participants were then given feedback as to the amount won on the trial. The maximal amount that could be won at the end of the experiment was 50 Swiss francs. Two training sessions (10 trials each) were conducted – one outside and one inside the scanner. The experimental session consisted of two runs, 36 trials each. The duration of every trial was ∼10s: cue (0.75s), ISI (2.5 – 3s), target (1s max), outcome (2s). The inter-trial interval was jittered from 1 to 9 s with a mean of 3.5s. Each run lasted about 6 minutes.

### Functional imaging acquisition

Imaging data were collected using a Philips Achieva 3.0 T magnetic resonance scanner with a 32-channel SENSE head coil at the MR Centre of the Psychiatric Hospital, University of Zurich. Functional MRI scans were acquired in 2 runs with 195 images in each run. A gradient-echo T2*weighted echo-planar image (EPI) sequence with 38 slices acquired in ascending order was used. Acquired in-plane resolution was 3 × 3 mm^2^, 3 mm slice thickness and 0.5 mm gap width over a field of view of 240 × 240 mm, repetition time 2000 ms, echo time 25 ms and flip angle 82°. Slices were aligned with the anterior–posterior commissure. Anatomic data were acquired using an ultrafast gradient echo T1-weighted sequence in 160 sagittal plane slices of 240 × 240 mm resulting in 1 × 1 × 1 mm voxels.

### Data analysis

Data analysis followed the same pipeline as in our previous work [20].

Functional magnetic resonance imaging data was analysed using SPM8 (Statistical Parametric Mapping, Wellcome Department of Cognitive Neurology, London, UK). Statistical analyses were performed using R (R Core Team, 2020).

### Behavioural data analysis

Reaction times to the target were the main outcome measure of the MID task. A two-way repeated measures ANOVA was conducted on RTs, with group as between subject factor and the reward condition (no, small and large reward) as within subject factor. Bonferroni-corrected pairwise comparisons were calculated as post-hoc tests for significant main effects. One BD-I subject was excluded only from the behavioural analysis due to corrupted data files.

### Image preprocessing

Data was preprocessed as described in Kirschner et al. 2019, 2016a, 2016b (see Supplemental material). To assure adequate quality of fMRI data motion and susceptibility artifacts were detected using the Art toolbox (http://web.mit.edu/swg/software.htm) and detected outlier scans were replaced by the mean image of the session for each participant. No subjects were removed from the analysis as a result of this procedure.

### First-level image analyses

Following Kirschner and colleagues (2018), in a first step, we used a general linear model with a parametric design to identify brain regions encoding the amount of reward obtained during the outcome phase. For this, each reward outcome condition (no-, small- and large reward) was modelled separately. The three outcome regressors accounted for the mean activations in the three conditions and allowed us to assess potential effects of group on small (0 – 0.40 CHF) and large (0 – 2 CHF) reward outcomes in general. Importantly, the small and large outcome regressors were parametrically modulated by the outcome received by each participant in each trial during the experiment (pmod small reward and pmod large reward respectively). The two parametric modulators capture linear deviations from the mean activity induced by the trial-specific reward level and are orthogonal to the mean regressors. The following regressors of no interest were used: one regressor for the anticipation phase (duration between 3.25 and 3.75 s), one regressor for target presentation, and, finally, one regressor for trials where participants made an error (modelled at target presentation). Thus, eight regressors were used for the first level analysis. All the explanatory variables were convolved using the canonical haemodynamic response function. We note that the two parametrically modulated reward regressors of interest were not correlated with the anticipation regressor, the latter serving to account for unspecific visual activations caused by stimulus presentation during the anticipation phase.

### Second-level image analyses

At the second (i.e. group) level, we interrogated the individual contrast images from the first-level parametric modulators in a two-step procedure.

First, we identified brain regions processing the reward amount received in the outcome phase, by assessing the contrast (pmod small reward + pmod large reward) in a voxel-wise whole brain analysis across all participants with a one-tailed t-test. The statistical threshold was set to P < 0.05, whole-brain voxel-level family-wise error (FWE) rate corrected for multiple comparisons. Our previous work [19, 20] has shown deficient adaptive coding in the right caudate and right insula/right precentral gyrus of SZ patients. We thus chose these regions among those showing a significant effect in the above analysis and used them as regions of interest (ROIs) in the next step.

In the second step, we investigated adaptive coding of reward across the three groups. The following adaptive coding contrast was run within the ROIs identified in step 1 (i.e. right caudate and right precentral gyrus): the contrast estimates of the large reward parametric regressor were subtracted from the contrast estimates for the small reward parametric regressor (pmod small reward – pmod large reward). A one-tailed voxel-wise t-test was used.

To investigate between-group differences, we extracted the mean contrast estimates of the adaptive coding contrast obtained in step 2 (pmod small reward – pmod large reward) from the reward sensitive regions obtained in step 1 (pmod low reward + pmod high reward) with the Marsbar toolbox. We then compared the three groups using Fischer’s one-way ANOVAs for each region and Tukey post-hoc comparisons.

To test for any associations between different symptoms and the adaptive coding contrast estimates in the BD group, we performed two-tailed Spearman rank correlation or Pearson correlation analyses within the regions showing significant group differences.

## Results

### Demographic and clinical data

Group characteristics are summarised in Table1. Compared to SZ participants, BD participants were older, had longer illness duration, more years of education, as well as a higher level of functioning (as indicated by higher GAF and PSP scores). 18 out of 25 BD participants were treated with atypical antipsychotics; 18 participants were prescribed mood stabilizers and 7 participants received antidepressants. All SZ participants were treated with atypical antipsychotics. Mean chlorpromazine equivalents were lower in the BD group compared to the SZ group (BD-I: 185.99 ± 259.8, SZ: 508.01 ± 369.2; U = 133, p < 0.001). None of the patients received typical antipsychotics.

### Behavioural results

Response time (RT) was the main measure of task performance. An ANOVA on the RTs in the three conditions showed no significant effect of group (F(2, 73)=1.54, p=0.22) nor condition * group interaction (F(4,142)=1.92, p=0.11), but, as in our previous studies, a significant effect of reward (F(2,73)=65.4, p<0.001). Specifically, larger rewards were associated with faster RTs. Furthermore, one way ANOVAs revealed no group differences for total accuracy (F(2, 47.1)=2.25, p=0.12) but for total amount won during the task (F(2, 48.2)=4.09, p=0.023). Tukey post-hoc tests showed that participants with BD-I won more than participants with SZ (p=0.03) and the amounts won did not differ between BD-I and HC (p=0.9). Overall, these results show that all groups were able to perform the task.

### Adaptive coding of reward: Group analysis

In the first step, brain areas processing reward amount were identified in a voxel-wise whole brain analysis across the three groups of participants (cluster defining threshold p=0.0005, FWE peak-level correction p<0.05). The identified regions showed increased activation with increasing reward during the outcome phase of the experiment. The results mirror the activations reported in our previous work (Table 2), as well as other studies on reward processing, and show significant effects in the right caudate and the right precentral region. Based on our previous work showing deficient adaptive coding in SZ in these regions, they were chosen as ROIs for step 2.

**Table 2.** Whole brain analyses of reward coding regions across all participants

|  | X | Y | Z | Cluster size | t |
| --- | --- | --- | --- | --- | --- |
| Right precentral gyrus | 60 | 3 | 12 | 250 | 6.77 |
|  | 55.5 | -3 | 7.5 |  | 4 |
| Right posterior caudate | 21 | -4.5 | 27 | 574 | 5.43 |
|  | 19.5 | -15 | 24 |  | 5.30 |
|  | 25.5 | -31.5 | 28.5 |  | 5.26 |
| Right anterior caudate | 16.5 | 19.5 | 16.5 | 145 | 5.12 |
| Left middle frontal gyrus | -24 | 22.5 | 54 | 2957 | 6.65 |
|  | -16.5 | 18 | 61.5 |  | 6.28 |
|  | -16.5 | 33 | 54 |  | 5.95 |
| Left angular gyrus | -48 | -60 | 40.5 | 1814 | 6.09 |
|  | -31.5 | -61.5 | 24 |  | 5.15 |
|  | -57 | -39 | 42 |  | 4.96 |
| mOFC | -7.5 | 54 | -6 | 456 | 6.08 |
|  | -1.5 | 63 | -1.5 |  | 4.79 |
|  | -1.5 | 57 | 9 |  | 4.04 |
| Right superior occipital gyrus | 15 | -88.5 | 24 | 1582 | 6.00 |
|  | 15 | -93 | 13.5 |  | 5.72 |
|  | 3 | -75 | 22.5 |  | 5.17 |
| Left orbitofrontal cortex | -9 | 24 | -16.5 | 1443 | 5.75 |
|  | -19.5 | 4.5 | -12 |  | 5.68 |
|  | -13.5 | 9 | -18 |  | 5.35 |
| Left postcentral gyrus | -21 | -31.5 | 61.5 | 406 | 5.15 |
|  | -21 | -33 | 76.5 |  | 4.52 |
|  | -28.5 | -28.5 | 66 |  | 4.19 |
| Right postcentral gyrus | 28.5 | -40.5 | 63 | 301 | 5.13 |
|  | 27 | -45 | 72 |  | 4.44 |
|  | 31.5 | -39 | 55.5 |  | 4.42 |
| Left caudate | -19.5 | -3 | 27 | 471 | 5.49 |
|  | -18 | 6 | 24 |  | 4.43 |
|  | -21 | 13.5 | 22.5 |  | 4.3 |
| Right middle frontal gyrus | 30 | 33 | 13.5 | 486 | 5.04 |
|  | 9 | 30 | 4.5 |  | 4.56 |
|  | -3 | 27 | 6 |  | 4.56 |
| Right putamen | 22.5 | 6 | -10.5 | 492 | 4.95 |
|  | 22.5 | 25.5 | -6 |  | 4.72 |
|  | 24 | 16.5 | -9 |  | 4.48 |
| Left posterior ventral cingulate cortex | -4.5 | -49.5 | 31.5 | 977 | 4.72 |
|  | 0 | -58.5 | 25.5 |  | 4.42 |
|  | -4.5 | -55.5 | 16.5 |  | 4.18 |

In step 2, we compared the three groups in a single analysis. A one-tailed t-test with the adaptive coding contrast in the three groups was run in the ROIs identified in step 1, showing a network of regions responding to reward adaptively (Table 3).

**Table 3.** Analysis of adaptive coding of reward across all participants

|  | X | Y | Z | Cluster size | t |
| --- | --- | --- | --- | --- | --- |
| Right precentral gyrus | 60 | 3 | 12 | 175 | 5.99 |
| Right caudate | 19.5 | -15 | 24 | 157 | 4.48 |
|  | 21 | -6 | 30 |  | 3.97 |
|  | 19.5 | 3 | 22.5 |  | 3.70 |
| Left postcentral gyrus | -19.5 | -31.5 | 60 | 152 | 5.14 |
|  | -22.5 | -30 | 67.5 |  | 3.47 |
| Left precentral gyrus | -33 | -15 | 55.5 | 107 | 4.85 |
| Left middle fontal gyrus | -24 | 22.5 | 54 | 221 | 4.59 |
|  | -16.5 | 33 | 54 |  | 4.52 |
| Right paracentral lobule | 4.5 | -27 | 61.5 | 157 | 4.46 |
|  | 3 | -39 | 60 |  | 3.62 |
| Right superior occipital gyrus | 15 | -88.5 | 22.5 | 159 | 4.25 |
|  | 16.5 | -94.5 | 9 |  | 4.15 |
|  | 22.5 | -90 | 12 |  | 3.40 |
| Left caudate | -21 | -6 | 25.5 | 144 | 4.06 |
|  | -18 | 6 | 24 |  | 3.58 |

To test for group differences in adaptive coding, we extracted activations for the adaptive coding contrast from the right caudate and the right precentral/insula region using the Marsbar toolbox and compared them across groups using Fisher’s one-way ANOVAs. Two regions of the right caudate showed a significant group difference (Figure 2). In the more anterior region (F(2,73)=6.03, p=0.004) the SZ group showed significantly weaker adaptive coding than the HC (p=0.02) and BD groups (p=0.007), while the BD group was at the same level as HC (p=0.9). In the more posterior region (F(2,73)=5.1, p=0.008), the SZ group was not significantly lower than the HC group (p=0.072), but the BD group was (p=0.008). There was no significant difference between the two patient groups (p=0.6). Thus, while patient with SZ show impaired adaptive coding in both caudate subregions, BD patients showed impaired adaptive coding only in the posterior caudate.

**Figure 2.**
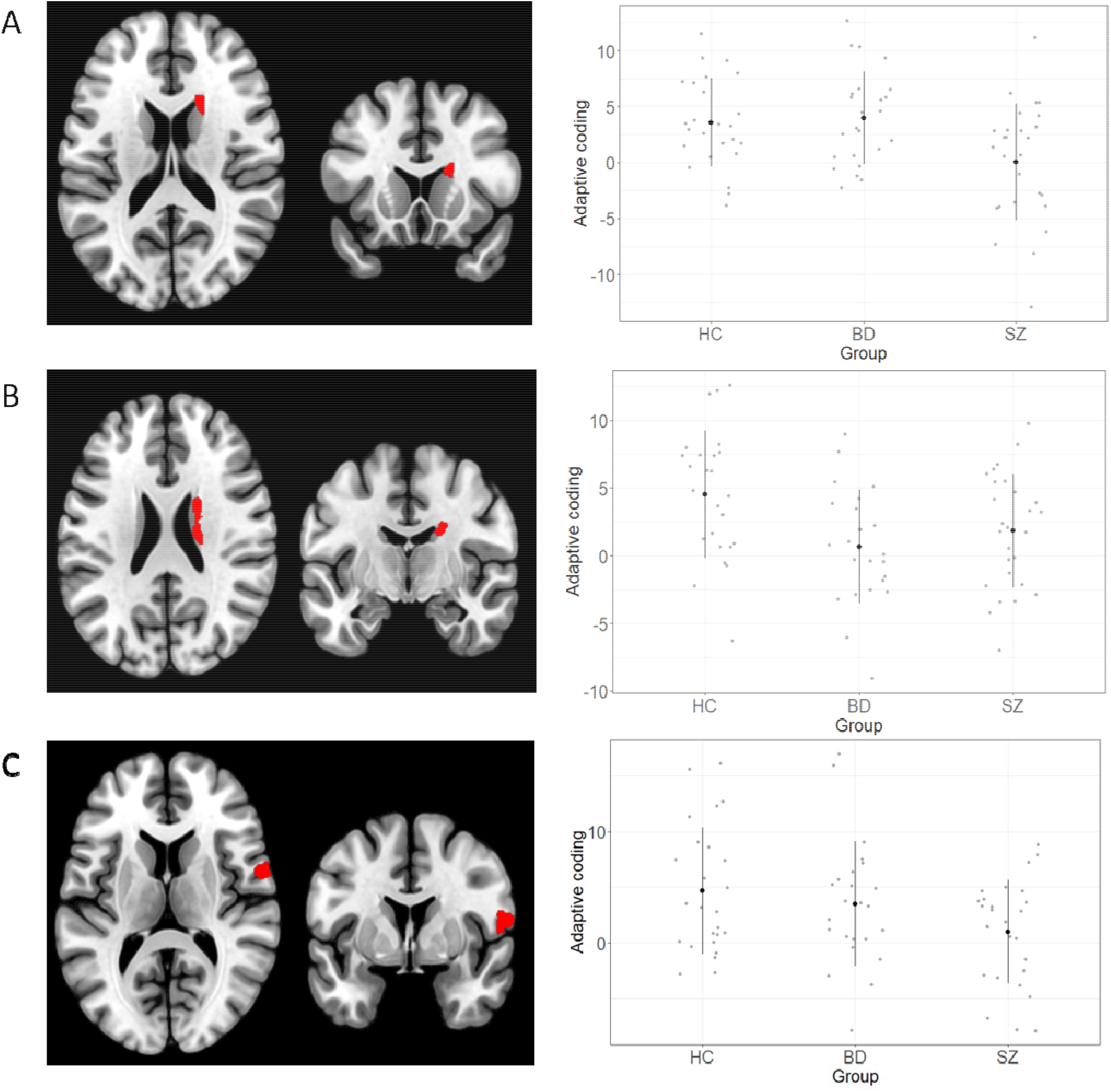
Reward-sensitive regions showing group differences in adaptive coding. Left panels: reward sensitive regions ((pmod small reward)+(pmod large reward) used as ROI for the adaptive coding contrast ((pmod small reward)-(pmod large reward; right panels). A: anterior right caudate. B: more posterior right caudate. C: right precentral gyrus.

In addition, a significant group difference was also observed also for the right precentral gyrus BA 6/insula (F(2,73)=3.3, p=0.043). As in our previous results, the SZ group showed significantly weaker adaptive coding compared to the HC group (p=0.038). Patients with BD, on the other hand, were not different from either the HC group (p=0.7), or the SZ group (p=0.2). This suggests that in the precentral/insular region adaptive coding in BD patients is situated between HC and patients with SZ.

### Correlational analyses in the BD-I group

In BD, adaptive coding in the more anterior part of the right caudate correlated positively with the total amount won in the task (r_p_=0.48 p=0.017).

In terms of symptomatology, a significant negative correlation was observed between adaptive coding in the more posterior section of the right caudate and the general subscale of the PANSS (r_s_= -0.45 p=0.029). No correlations between adaptive coding and other psychopathology measures (HAMD, CSDS, YMRS, PANSS positive or total subscales) were found.

Additionally, a trend-level positive correlation was observed between adaptive coding in the right precentral gyrus and chlorpromazine equivalents for antipsychotic medication (r_s_=0.38, p=0.065).

## Discussion

The present work explored adaptive coding in patients with BD-I using a version of the monetary incentive delay task, comparing their brain response to rewards to that of patients with SZ and HC. As in SZ, adaptive coding appears to be impaired in BD-I. However, we observed several differences between SZ and BD patients. Specifically, adaptive coding was significantly reduced in BD-I participants in the more posterior part of the right caudate but preserved in its more anterior part, in contrast to patients with SZ. In the right precentral gyrus, BD-I patients showed intermediate levels of neural adaptation to reward.

Similar to patients with SZ, BD-I patients showed adaptive coding deficits in the more posterior part of the right caudate, a region that underlies reward-guided action selection and reward learning. In BD, reduced activation in the right caudate during reward feedback has previously been reported in a card-guessing paradigm[10] for patients who were experiencing a depressive episode. Our previous work in SZ participants has shown adaptive coding in the caudate to be negatively associated with increased total symptom severity, as measured by the PANSS, but also its general and negative subscales. In BD patients, we observed a negative association between adaptive coding in the posterior part of the right caudate (where deficient adaptive coding was observed in this group) and the PANSS general subscale. Thus, in both disorders adaptive coding may represent a general deficit, spanning multiple areas of motivation, perception and cognition. Indeed, dopamine neurons send widespread projections across the brain and their insensitivity to context might underlie a broad range of deficits. In BD, no associations were found between specifically depressive (as measured by the HAMD), manic or psychotic symptoms and adaptive coding.

Interestingly, in the BD group, adaptive coding in the more anterior part of the right caudate (where adaptive coding appeared intact in this group) correlated positively with the amount won on the task, i.e. the stronger the adaptive coding, the more participants won. Thus, one could speculate that to the extent adaptive coding occurred in the anterior right caudate it rescued performance. However, this speculation is to be taken with caution due to the small sample size and small differences between groups. Moreover, our findings seem to imply a segregation of the adaptive coding deficit between BD and SZ in the right caudate. Potentially, this deficit segregation between the two conditions might relate to a functional segregation of the caudate. Previous work in healthy participants has indeed indicated that the more anterior part of the caudate is engaged in learning new rules and more cognitive and emotional processing, whereas the more posterior part is dedicated to action-based processes [34-37].

As in our previous work, SZ patients showed an adaptive coding deficit in the right precentral gyrus/insula. BD participants showed no difference from HC in this region. However, they were also not significantly different from SZ participants, in line with intermediate levels of adaptive coding. In addition to the functional results described above, previous anatomical studies have shown a decrease in grey matter volume in BD patients in the right caudate and right as well as left precentral gyri [38, 39]. Of note, adaptive coding in the right precentral gyrus in BD positively correlated at trend level with the dose of antipsychotic medication they received. We can thus speculate that antipsychotic medication in BD may have a positive effect on context adaptation in this brain region [40].

Thus, it appears that both BD and SZ show deficient adaptive coding. Although this deficit is less pronounced in BD, it could still point towards a continuity in reward processing adaptive coding alterations between the two conditions. Similarly to schizophrenia, a blunted contextual response could be inherent to BD, and the difference between the two conditions could simply be quantitative. Deficient adaptive coding in BD would also be in line with previous research hinting at reduced contextual adaptation related to manic and hypomanic symptoms. Indeed, previous work in healthy participants prone to hypomania has shown a lack of discrimination between stimulus values [41], with neutral outcomes activating the medial temporal lobe similarly to rewards (while controls showed increased activation for rewards). In addition, an earlier study in BD patients performing the MID task during a manic episode found no difference in nucleus accumbens responses induced by receipt and omission of reward at the outcome phase of the task [11]. Thus, these earlier studies appear to point towards a deficit in contextual adaptation in mania.

In contrast to our findings, some previous work has found no differences between BD patients and HC in reward processing at the outcome phases of different tasks [6, 13, 14, 42]. It is thus possible that adaptive coding represents a more fine-grained assessment of reward processing, beyond simple evaluation of brain activity during the outcome phase of reward tasks. It should also be noted that no significant differences have been observed in brain activity during reward anticipation between the three groups analysed here [3]. Thus, despite similar behavioural performance and reward anticipation in HC and BD, the latter could still harbour a deficit in adaptation to reward context.

It is also possible that adaptive coding deficits have a different aetiology in BD and SZ. Limitations to transdiagnostic approaches have indeed been voiced, highlighting the fact that, for reward in particular, different mechanisms could lead to similarly altered brain activity patterns (“equifinality”) [43, 44]. In SZ, an adaptive response to reward would be prevented by a simultaneous decrease of adaptive dopamine transients and an increase in spontaneous dopamine transients in the striatum. This would result in a blunted response to rewarding stimuli as well as an increased response to neutral cues [45], thus reducing the neural discrimination between the two. In contrast, in BD, the substantial changes in internal state (manic vs depressive) may lead to a blunted processing of external contexts [41]. Although the exact role of dopamine in BD is still a matter of debate, previous work points towards an increase of striatal dopaminergic transmission during mania, and a decrease in the dopaminergic function during depression [46, 47]. Such extreme fluctuations of the dopaminergic response could lead to its overall blunting, which in turn may result in reduced discriminability of reward stimuli in euthymia, mania and depression. This hypothesis is in line with the interpretation of Redlich and colleagues (2015), who found significantly reduced caudate activation in depressed BD patients compared to patients with unipolar depression (UD). They concluded that BD patients might exhibit a relatively stronger impairment of the mesolimbic structures because they have to regulate the change between manic/hypomanic and depressive states.

Several limitations to the present work should be pointed out. A larger sample size would have allowed to draw more robust conclusions, especially for the right precentral gyrus, where BD patients did not differ significantly from either of the other two groups. Our sample of euthymic patients also included patients with sub-syndromal depressive symptoms [22]. Thus, we did not use a conservative definition of euthymia [48].

In conclusion, we demonstrate for the first time that patients with BD-I disorder show adaptive coding deficits, similar to those observed in SZ patients. Adaptive coding in BD appeared more preserved as compared to SZ participants especially in the more anterior part of the right caudate and to a lesser extent also in the right precentral gyrus. These results reinforce the importance of context processing and adaptive coding deficits across mental illnesses [49, 50]. Future work will establish whether similar mechanisms are involved in context processing deficits across different disorders.

## Acknowledgements

The work reported here was funded by the Swiss National Science Foundation (Grants No. <u>105314_140351 and 10001CL_169783 to SK</u>).

## Disclosures

All authors report no biomedical financial interests or potential conflicts of interest

## Notes

### Competing Interest Statement

The authors have declared no competing interest.

## References

1. Yamada, Y., et al., Specificity and continuity of schizophrenia and bipolar disorder: relation to biomarkers. Current pharmaceutical design, 2020. 26(2): p. 191.

2. Kirschner, M., et al., From apathy to addiction: Insights from neurology and psychiatry. Progress in Neuro-Psychopharmacology and Biological Psychiatry, 2020. 101: p. 109926.

3. Kirschner, M., et al., Shared and dissociable features of apathy and reward system dysfunction in bipolar I disorder and schizophrenia. Psychological medicine, 2020. 50(6): p. 936–947.

4. Schreiter, S., et al., Neural alterations of fronto-striatal circuitry during reward anticipation in euthymic bipolar disorder. Psychological Medicine, 2016. 46(15): p. 3187–3198.

5. Nielsen, M.Ø., et al., Alterations of the brain reward system in antipsychotic naïve schizophrenia patients. Biological psychiatry, 2012. 71(10): p. 898–905.

6. Nusslock, R., et al., Waiting to win: elevated striatal and orbitofrontal cortical activity during reward anticipation in euthymic bipolar disorder adults. Bipolar disorders, 2012. 14(3): p. 249–260.

7. Mason, L., et al., Decision-making and trait impulsivity in bipolar disorder are associated with reduced prefrontal regulation of striatal reward valuation. Brain, 2014. 137(8): p. 2346–2355.

8. Dutra, S.J., et al., Elevated striatal reactivity across monetary and social rewards in bipolar I disorder. Journal of abnormal psychology, 2015. 124(4): p. 890.

9. Trost, S., et al., Disturbed anterior prefrontal control of the mesolimbic reward system and increased impulsivity in bipolar disorder. Neuropsychopharmacology, 2014. 39(8): p. 1914–1923.

10. Redlich, R., et al., Reward processing in unipolar and bipolar depression: a functional MRI study. Neuropsychopharmacology, 2015. 40(11): p. 2623–2631.

11. Abler, B., et al., Abnormal reward system activation in mania. Neuropsychopharmacology, 2008. 33(9): p. 2217–2227.

12. Schwarz, K., et al., Transdiagnostic prediction of affective, cognitive, and social function through brain reward anticipation in schizophrenia, bipolar disorder, major depression, and autism spectrum diagnoses. Schizophrenia bulletin, 2020. 46(3): p. 592–602.

13. Chase, H.W., et al., Dissociable patterns of abnormal frontal cortical activation during anticipation of an uncertain reward or loss in bipolar versus major depression. Bipolar disorders, 2013. 15(8): p. 839–854.

14. Smucny, J., et al., Schizophrenia and bipolar disorder are associated with opposite brain reward anticipation-associated response. Neuropsychopharmacology, 2021. 46(6): p. 1152–1160.

15. Smirnakis, S.M., et al., Adaptation of retinal processing to image contrast and spatial scale. Nature, 1997. 386(6620): p. 69–73.

16. Kourtzi, Z. and C.E. Connor, Neural representations for object perception: structure, category, and adaptive coding. Annual review of neuroscience, 2011. 34: p. 45–67.

17. Tobler, P.N., C.D. Fiorillo, and W. Schultz, Adaptive coding of reward value by dopamine neurons. Science, 2005. 307(5715): p. 1642–1645.

18. Burke, C.J., et al., Partial adaptation of obtained and observed value signals preserves information about gains and losses. Journal of Neuroscience, 2016. 36(39): p. 10016–10025.

19. Kirschner, M., et al., Deficits in context-dependent adaptive coding of reward in schizophrenia. NPJ schizophrenia, 2016. 2(1): p. 1–8.

20. Kirschner, M., et al., Deficits in context-dependent adaptive coding in early psychosis and healthy individuals with schizotypal personality traits. Brain, 2018. 141(9): p. 2806–2819.

21. Jauhar, S., et al., A test of the transdiagnostic dopamine hypothesis of psychosis using positron emission tomographic imaging in bipolar affective disorder and schizophrenia. JAMA psychiatry, 2017. 74(12): p. 1206–1213.

22. Tohen, M., et al., The International Society for Bipolar Disorders (ISBD) Task Force report on the nomenclature of course and outcome in bipolar disorders. Bipolar disorders, 2009. 11(5): p. 453–473.

23. Young, R.C., et al., A rating scale for mania: reliability, validity and sensitivity. The British journal of psychiatry, 1978. 133(5): p. 429–435.

24. Hamilton, M., Development of a rating scale for primary depressive illness. British journal of social and clinical psychology, 1967. 6(4): p. 278–296.

25. Addington, D., J. Addington, and B. Schissel, A depression rating scale for schizophrenics. Schizophrenia research, 1990. 3(4): p. 247–251.

26. Kirkpatrick, B., et al., The brief negative symptom scale: psychometric properties. Schizophrenia bulletin, 2011. 37(2): p. 300–305.

27. Mucci, A., et al., The Brief Negative Symptom Scale (BNSS): independent validation in a large sample of Italian patients with schizophrenia. European Psychiatry, 2015. 30(5): p. 641–647.

28. Frances, A., H. Pincus, and M. First, The global assessment of functioning scale (GAF). Diagnostic and Statistical Manual of Mental Disorders, 1994. 4.

29. Juckel, G., et al., Validation of the Personal and Social Performance (PSP) Scale in a German sample of acutely ill patients with schizophrenia. Schizophrenia research, 2008. 104(1-3): p. 287–293.

30. Kay, S.R., L.A. Opler, and J.-P. Lindenmayer, The positive and negative syndrome scale (PANSS): rationale and standardisation. The British Journal of Psychiatry, 1989. 155(S7): p. 59–65.

31. Fervaha, G., et al., Examination of the validity of the Brief Neurocognitive Assessment (BNA) for schizophrenia. Schizophrenia research, 2015. 166(1-3): p. 304–309.

32. Simon, J.J., et al., Reward system dysfunction as a neural substrate of symptom expression across the general population and patients with schizophrenia. Schizophrenia bulletin, 2015. 41(6): p. 1370–1378.

33. Cools, R., et al., Tryptophan depletion disrupts the motivational guidance of goal-directed behavior as a function of trait impulsivity. Neuropsychopharmacology, 2005. 30(7): p. 1362–1373.

34. Hampshire, A., et al., Probing cortical and sub-cortical contributions to instruction-based learning: Regional specialisation and global network dynamics. NeuroImage, 2019. 192: p. 88–100.

35. Brovelli, A., et al., Differential roles of caudate nucleus and putamen during instrumental learning. Neuroimage, 2011. 57(4): p. 1580–1590.

36. Robinson, J.L., et al., The functional connectivity of the human caudate: an application of meta-analytic connectivity modeling with behavioral filtering. Neuroimage, 2012. 60(1): p. 117–129.

37. Mattfeld, A.T. and C.E. Stark, Striatal and medial temporal lobe functional interactions during visuomotor associative learning. Cerebral Cortex, 2011. 21(3): p. 647–658.

38. Lisy, M.E., et al., Progressive neurostructural changes in adolescent and adult patients with bipolar disorder. Bipolar disorders, 2011. 13(4): p. 396–405.

39. Lyoo, I.K., et al., Frontal lobe gray matter density decreases in bipolar I disorder. Biological psychiatry, 2004. 55(6): p. 648–651.

40. Liemburg, E.J., et al., Antipsychotic medication and prefrontal cortex activation: a review of neuroimaging findings. European Neuropsychopharmacology, 2012. 22(6): p. 387–400.

41. O’Sullivan, N., et al., fMRI evidence of a relationship between hypomania and both increased goal-sensitivity and positive outcome-expectancy bias. Neuropsychologia, 2011. 49(10): p. 2825–2835.

42. Johnson, S.L., et al., Neural responses to monetary incentives in bipolar disorder. NeuroImage: Clinical, 2019. 24: p. 102018.

43. Whitton, A.E., M.T. Treadway, and D.A. Pizzagalli, Reward processing dysfunction in major depression, bipolar disorder and schizophrenia. Current opinion in psychiatry, 2015. 28(1): p. 7.

44. Weinberger, D.R. and T.E. Goldberg, RDoCs redux. World Psychiatry, 2014. 13(1): p. 36.

45. Maia, T.V. and M.J. Frank, An integrative perspective on the role of dopamine in schizophrenia. Biological psychiatry, 2017. 81(1): p. 52–66.

46. Cousins, D.A., K. Butts, and A.H. Young, The role of dopamine in bipolar disorder. Bipolar disorders, 2009. 11(8): p. 787–806.

47. Ashok, A.H., et al., The dopamine hypothesis of bipolar affective disorder: the state of the art and implications for treatment. Molecular psychiatry, 2017. 22(5): p. 666–679.

48. Samalin, L., et al., Course of residual symptoms according to the duration of euthymia in remitted bipolar patients. Acta Psychiatrica Scandinavica, 2016. 134(1): p. 57–64.

49. Northoff, G. and H. Mushiake, Why context matters? Divisive normalization and canonical microcircuits in psychiatric disorders. Neuroscience research, 2020. 156: p. 130–140.

50. Rosenberg, A. and A. Sunkara, Does attenuated divisive normalization affect gaze processing in autism spectrum disorder? A commentary on Palmer et al.(2018). Cortex, 2019. 111: p. 316–318.

