## Supplementary material for "Comparing adaptive coding of reward in bipolar I disorder and schizophrenia"

**Supplemental material**

**Acquisition parameters**

Imaging data was collected with a Philips Achieva 3.0T magnetic resonance (MR) scanner using a 32 channel SENSE head coil (Philips, Best, The Netherlands) at the Psychiatric Hospital of the University of Zurich. Functional MRI (fMRI) was acquired in two runs with 195 ascending transverse plane images using a gradient-echo T2*-weighted echo-planar image (EPI) sequence over the whole brain. Acquired in-plane resolution was 3×3mm2, 3mm slice thickness and 0.5mm gap width over a field of view of 240×240mm2, a repetition/echo time (TR/TE) of 2000/25ms and a flip angle of 82°. The first five scans were discarded to account for T1 saturation effects. Slices were aligned with the anterior–posterior commissure. Anatomical data was acquired with an ultrafast gradient echo T1-weighted sequence in 160 sagittal plane slices of 240×240mm2 resulting in 1x1x1mm3 voxels.

**Image preprocessing**

Functional images were corrected for differences in the time of slice acquisition. The Realign and Unwarp functions of SPM8 were used to correct our data for head motion, with an allowed translational head motion limited to ±4 mm. A voxel displacement map, calculated from double phase and magnitude field map data, was used to correct for combined static and dynamic distortions. We performed segmentation, bias correction, and spatial normalization. Finally, images were smoothed using a Gaussian kernel of 6mm width at half-maximum.
